# The reduced prevalence of macrolide resistance in *Mycoplasma pneumoniae* clinical isolates from pediatric patients in Beijing in 2016

**DOI:** 10.1101/339317

**Authors:** Xiujun Tian, Ran Wei, Junyan Shao, Hong Wang, Jing Li, Wei Zhou, Xuanguang Qin, Yinghui Hu, Haiwei Dou, Dongxing Guo, Jingyi Li, Dan Li, Baoping Xu, Deli Xin

## Abstract

Older children especially from seven to thirteen years old are more prone to develop *Mycoplasma pneumoniae* (MP) infection; in winter children are more susceptible to infect with MP. In Beijing, China in 2016 the rates of macrolide resistance of MP were 69.48% (in total children), 61.59% (in outpatients) and 79.28% (in hospitalized patients), respectively. All the macrolide resistant isolates harbored A2063G or A2064G mutation in the 23S rRNA gene. Seven isolates showed a mixed infection. Susceptibility results showed that 73 isolates with the A2063G mutation demonstrated different levels resistance to erythromycin (MIC=8 to>256μg/ml), azithromycin (MIC=8 to>64μg/ml) and josamycin (MIC=2 to 8μg/ml). No cross-resistance was observed in the in the antibiotics of levofloxacin and tetracycline against MP.

---

*Mycoplasma pneumoniae* (MP) is an important leading cause of community-acquired pneumonia (CAP) in children, accounting for 10%–40% of CAP cases^1^. Macrolide-resistant MP (MRMP) was first isolated in Asia and has rapidly increased over the past decades^2^. Our previous study found that 46 clinical isolates collected from 2003 to 2006 were MRMP, the rate of macrolide resistance was as high as 92% in Beijing^3^. Macrolide resistance rates were 3.5-13.2% in USA^4–5^, below 10% in Europe^6^, 50-90% in Japan^7–8^ and more than 90% in China^3,9–10^.

Tanaka et al found that the prevalence of macrolide-resistant MP gradually declined during 2013–2015^11^. In China few studies about macrolide-resistant were reported since 2015, this study was scheduled to investigate the epidemiology and macrolide resistance rate of pediatric patients with MP infection in Beijing in 2016.

## MATERIALS AND METHODS

### *M. pneumoniae* strains

A total of 619 pediatric patients suspected as MP infection at seven hospitals in Beijing (China Meitan General Hospital, Civil Activation General Hospital, Beijing Changping Hospital of Integrated Chinese and Western Medicine, Peking University Third Hospital, Beijing Chao-Yang Hospital, New Century International Children’s Hospital, and Beijing Friendship Hospital) were enrolled in the study. During January 2016 to December 2016, patients aged from 1 month to 14 years, had a preceding fever, cough or pharyngalgia whose onset time was from 1 to 7 days were eligible for enrollment. Throat swabs or BALF were obtained in 24h after their enrollment.

Positive result for culture or DNA detection by real-time PCR from throat swabs or BALF was considered as MP infection. For MP culture, clinical specimens were vortexed, supplemented with amphotericin B and penicillin, and inoculated into SP-4 medium. The medium was incubated at 37°C, and observed daily for 2–6 weeks for a decrease in pH (a red to yellow color change). Then the positive specimens for MP culture were identified using a real-time PCR^12^. DNA of MP in throat swabs or BALF was extracted using a QIAmp DNA Mini Kit (Qiagen, Hilden, Germany).Then, the above real-time PCR was done to detect DNA of MP.

### Detection of the mutations associated with macrolide resistance of MP

Macrolide resistance associated mutations in domain V of the 23S rRNA gene were detected using a direct sequencing method as previously reported^13^. The amplification product of each MP strain, including the reference strain M129 (ATCC 29342) was sequenced by the Invitrogen Biotechnology Co., Ltd.

### Antimicrobial susceptibility testing

The minimum inhibitory concentrations (MICs) of five macrolides (erythromycin, azithromycin, levofloxacin, tetracycline and josamycin) for MP strains were determined by broth microdilution methods with SP4 broth (Remel). MP reference strain M129 was tested as an antibiotic-sensitive control. Susceptibility tests were performed in triplicate.

### Statistical analysis

All data was expressed as means and standard deviations, unless otherwise indicated. Differences in categorical variables were assessed with the χ2 test or Fisher’s exact test. All analyses were performed using SPSS for Windows, version 17.0 software (SPSS Inc., Chicago, IL, USA), and a two-sided P value <0.05 were considered statistically significant.

## RESULTS

### Epidemiology of pediatric patients with MP infection

A total of 262 (42.33%) pediatric patients were diagnosed as MP infection by culture or DNA detection of MP. Among the 238 patients with MP infection marked with age, the age ranged from 1.5 months to 14 years with a median of 5.67 years, Fifty-seven (23.95%) cases were 0-3 years, 71 (29.83%) were 4-6 years and 110 (46.22%) were 7-13 years. The detection rates of MP infection in the above different age groups were 52.38%, 34.13% and 34.13%, respectively. The incidence of MP infection increased with age. The prevalence of MP infection in patients aged 7-13 especially 7-9 years was significantly higher than those in the other two groups (P< .001).Among the 246 patients with MP infection marked with sex, male cases are 123, female cases are 123, the ratio of male to female was 1:1. MP infection predominated in fall and winter, 77 cases (29.84%) in fall (from September through November) and 89 (34.50%) cases in winter (from December through February)were involved, respectively, compared with 34 cases (13.18%) in spring (from March through May), and 58 (22.48%) cases in summer (from June through August). The prevalence of MP infection was much higher during winter than that in spring in pediatric patients (P < .05). The age and season distributions of MP infection in pediatric patients were shown in Fig. 1 and Fig. 2. Macrolide resistance rates of pediatric patients with MP infection. A total of 249 MP clinical isolates whose gene sequence in domain V of the 23S rRNA gene were successfully determined, 173 were MRMP strains. The rate of macrolide resistance was 69.48% (173/249) in pediatric patients with MP infection in Beijing in 2016. There were 138 outpatients and 111 hospitalized patients in the 249 pediatric patients. The rates of macrolide resistance in outpatients and inpatients were 61.59% (85/138) and 78.38% (88/111), respectively. The summary of the rates of macrolide resistance was shown in Table 1. The rates of macrolide resistance in different age groups are follows: 0-3 years, 70.18% (40/57); 4-6 years, 53.52% (38/71); 7-13 years, 70.91% (78/110). The rates of macrolide resistance in different seasons are follows: spring, 67.74% (21/31); summer, 72.88% (43/59); fall, 67.12% (49/73) and winter, 69.41% (59/85).

**Fig. 1.**
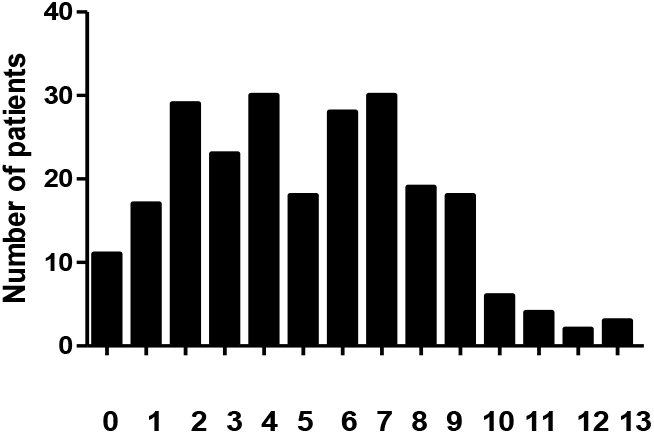
The age distribution of MP infection in pediatric patients

**Fig. 2.**
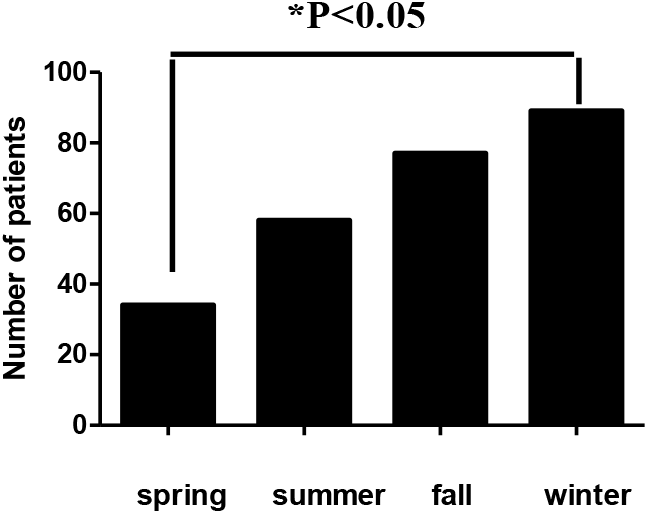
The season distribution of MP infection in pediatric patients

**Table 1.**
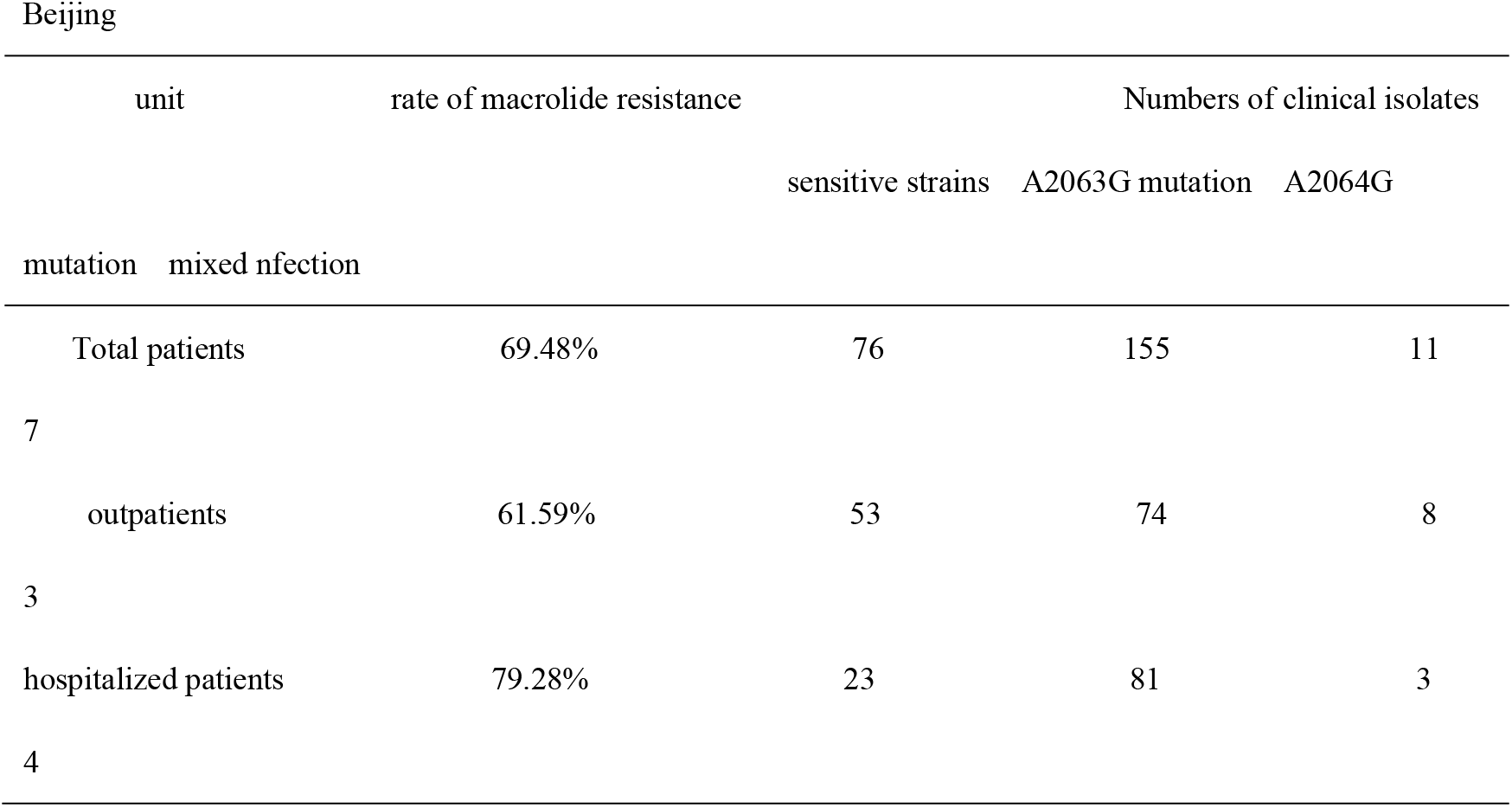
The rate of macrolide resistance in pediatric patients with MP infection in 2016 in Beijing

### Antimicrobial susceptibility testing

A total of 78 clinical isolates were randomly selected (7 isolates without domain V region mutations and 71 isolates with A2063G mutation) in the antimicrobial susceptibility testing. Compared with the results of 7 clinical isolates carrying a wild type of 23S rRNA gene and the MP reference strain M129, all the 71 clinical isolates with A2063G mutation except BS315 strain showed a high resistance to erythromycin (64 to>256μg/ml). The erythromycin-resistant strains showed cross-resistance to azithromycin and josamycin. The MIC of azithromycin (10 isolates with MIC_90_ value of 32μg/ml) was lower than that of erythromycin. The MIC of josamycin was the lowest (2 to 8μg/ml) in the three macrolides. The MIC_90_ values of BS315 strain to erythromycin, azithromycin and josamycin were 8μg/ml, 16μg/ml and 4μg/ml, respectively.

All the selected MP clinical isolates as well as the MP reference strain M129 were susceptible to levofloxacin and tetracycline. Levofloxacin and tetracycline showed similar MIC_90_ distribution for MP, the range of MIC_90_ values of levofloxacin and tetracycline were between 0.125 and 1μg/ml (Table 2).

**Table 2.** MIC_90_ range of five antimicrobial agents against 78 *M. pneumoniae* clinical isolates and M129

| Isolates | No. of strains | erythromycin | azithromycin | josamycin | levofloxacin |
| --- | --- | --- | --- | --- | --- |
| tetracycline (μg/ml) |  |  |  |  |  |
| Clinical sensitive strains | 7 | 0.0018-0.03 | 0.0005-0.004 | 0.015-0.0625 | 0.125-0.5 |
| 0.125-0.5 |  |  |  |  |  |
| M129 reference stain | 1 | 0.032 | 0.008 | 0.063 | 0.5 |
| 0.125 |  |  |  |  |  |
| A2063G mutation strains | 71 | 8->256 | 8->64 | 2-8 | 0.25-1 |
| 0.125-1 |  |  |  |  |  |

## DISCUSSION

To our knowledge, this is the first study about the evaluation of macrolide –resistance rate in pediatric outpatients with MP infection in Beijing, China. The fact of high prevalence of MRMP clinical isolates in pediatric patients in China mostly come from the data of hospitalized patients. Outpatients are not easily achievable as many MP infections such as mild tracheobronchitis are often undiagnosed, the macrolide resistance rate of MP in outpatients especially in pediatric outpatients is still unclear. To investigate this and monitor the situation of macrolide resistance in pediatric patients with MP infection, the present study was designed.

The infection rate of MP was 42.33% (262/619) in pediatric patients with respiratory infections in the present study. The infection of MP is related to age and season. The infection of MP was mainly prevalent in 7-13 especially 7-9 years old pediatric patients in the present study, which was in accordance with the previous conclusion that the main burden of the infection is typically in preschool and school-age children ^14–15^.Moreover, the prevalence of MP infection in patients aged 7-13 was significantly higher than those in the other two groups (P< .001) in our study. The peak season of MP infection was winter in our study, and there was seasonal difference for the prevalence of MP infection, it was higher during winter than that in spring (P< .05). These findings were consistent with the reports that the epidemic seasons in north of China is winter but is summer and autumn in the south of China^8,16^.

Macrolides usually are used as the first-line therapeutic drug for MP infection in children. Since the isolation of the first MRMP strain, mcrolide resistance rate has been increasing worldwide. However, since 2015 the reports about macrolide resistance rate of MP are rare in the world, no report in China. So our present analysis is significant because it reported on the recent macrolide resistance rate of MP in Chinese children. The macrolide resistance rate was 69.48% in pediatric patients with MP infection in Beijing, China in 2016. The macrolide resistance rates recently published were 100% in children of Zhejiang province, China in 2014^10^, 87.2% in South Korean children in 2015^17^, 43.6% in Japanese children in 2015^11^, 47.1% in children of Hong-Kong in 2014^18^, 13.2% in American children through 2012 to 2014^5^, 9.3% in English children between 2014 and 2015^19^. Compared with the above data, the prevalence of MRMP clinical isolates among children in China has significantly decreased to 69.48% from 80-100% ^3,10,13^. The decrease of high rate of macrolide resistance might be partially attributable to the inclusion of pediatric outpatients. The macrolide resistance rate in pediatric outpatients with MP infection was only 61.59%. This is the first report about the evaluation of macrolide resistance rate in pediatric outpatients with MP infection in Beijing, China. Cao et al ever investigated the rate of macrolide resistance in outpatients, but the patients in his study were adults and adolescents (aged≥14 years) ^20^. Ishiguro et al showed that the rate of macrolide resistance in pediatric outpatients in Hokkaido, Japan was 44.3%. However, the sample size in his study was small, there were only 31 pediatric outpatients enrolled from December 1, 2012 to July 31, 2014^8^.This is also the first report about the lowest rate of macrolide resistance in MP clinical isolates in pediatric patients in Beijing since 2009.The rate of macrolide resistance in pediatric hospitalized patients was 79.28%, which was also lower than those in most reports in Asian countries. This might be due to the gradually reduced prevalence of macrolide resistant MP infection. Tanaka et al found that the prevalence of MRMP was high in Japan during 2008-2012, gradually declined during 2013–2015. They thought the rate of macrolide resistance might be affected by changes in the use of oral macrolide agents^11^. Possibly, the situation in our country was similar to Japan. Thankfully, the situation of macrolide reisitance of MP was not as serious as previously reported in China.

Although MRMP clinical isolates are prevalent in worldwide, to our knowledge, the resistance mechanisms are still uncertain, the point mutation in the specific locus of the 23S rRNA gene region, especially in loci 2063 and 2064 is most commonly proposed^21–23^.In the present study, A2063G transition predominated in pediatric patients with MP infection, which involved 161 cases (up to 93.06%). Meanwhile, eleven A2064G transition cases were identified. A point mutation in the loci 2063 and 2064 plays an important role in the macrolide resistance in our study, no other macrolide-resistant related point mutations were identified. In addition, seven mixed infection cases were identified. Cardinale et al reported the first case showing the detection of macrolide-resistance in MP not at admission but after 10 days of clarithromycin treatment ^24^, Suzuki et al confirmed his finding in 21 children infected with MP. All the MP clinical strains shifted from macrolide sensitive at beginning to macrolide resistant after 7-24 days treatment of clarithromycin or azithromycin^25^. The above mentioned findings support a hypothesis that the emergence of mixed type of macrolide-resistant strains is possible selected outgrowth during drug administration.

Based on the results of the susceptibility test, the A2063G transitions are responsible for high-level resistance to 14- and 15-member ring macrolides, such as erythromycin(8 to>256 μ g/ml) and azithromycin (8 to>64 μ g/ml) in M. pneumoniae. However, 16-member ring macrolides, such as josamycin demonstrated middle-level resistance, with MICs of 2 to 8 μ g/ml against clinical strains with the A2063G mutation. Cross-resistance was not observed in the levofloxacin and tetracycline groups. All the isolates including the strains with macrolide-resistance associated mutations remained susceptible to levofloxacin and tetracycline. To date, no levofloxacin or tetracycline resistant strains have been isolated from MP clinical specimens. This situation of high macrolide resistance for first-line treatment drugs had caused great difficulties in the clinical treatment of MP pneumonia, especially in pediatric infections. The data of our susceptibility tests demonstrated that all the MP isolates were sensitive to levofloxacin and tetracycline. This finding suggests that in the situation of patients with the refractory MP pneumonia, such antibiotics can act as alternative medicines for treating MP infection in adults, however, tetracycline and levofloxacin are not approved for use in children under 8 years old and under 18 years old, respectively, in China.

In summary, older children especially 7-13 years old are more prone to developing MP infection; children are more susceptible to infect with MP in winter. The infection rate of MP was 42.33% in pediatric patients in Beijing in 2016. The rates of macrolide resistance of MP were 69.48% (in total children), 61.59% (in outpatients) and 79.28% (in hospitalized patients), respectively in Beijing, China in 2016, this is the first study about the evaluation of macrolide –resistance rate in pediatric outpatients with MP infection in Beijing, China. A point mutation in the loci 2063 A-G predominated in 93.06% of MRMP clinical isolates in Beijing population in 2016.

Cross-resistance was not observed in the antibiotics of levofloxacin and tetracycline against *Mycoplasma pneumoniae*.

## ACKNOWLEDGMENTS

This study was supported by National Natural Science Foundation of China (grant reference number: 81271890) and Beijing Municipal Science and Technology Commission (grant reference number: Z171100001017081).We declare that the experiments performed and described here comply with the current laws of the People’s Republic of China.

